# A Portable Structural Analysis Library for Reaction Networks

**DOI:** 10.1101/245068

**Authors:** Yosef Bedaso, Frank T. Bergmann, Kiri Choi, Herbert M. Sauro

## Abstract

The topology of a reaction network can have a significant influence on the network’s dynamical properties. Such influences can include constraints on network flows and concentration changes or more insidiously result in the emergence of feedback loops. These effects are due entirely to mass constraints imposed by the network configuration and are important considerations before any dynamical analysis is made. Most established simulation software tools usually carry out some kind of structural analysis of a network before any attempt is made at dynamic simulation. In this paper we describe a portable software library, `libStructural`, that can carry out a variety of popular structural analyses that includes conservation analysis, flux dependency analysis and enumerating elementary modes. The library employs robust algorithms that allow it to be used on large networks with more than a two thousand nodes. The library accepts either a raw or fully labeled stoichiometry matrix or models written in SBML format. The software is written in standard C/C++ and comes with documentation and a test suite. The software is available for Windows, Mac OS X, and can be compiled easily on any Linux operating system. A language binding for Python is also available through the pip package manager making it trivial to install on any standard Python distribution. As a second example, we also create a new `libStructural` plugin for PathwayDesigner that allows solutions to be viewed graphically. The source code is licensed under the open source BSD license and is available on GitHub (https://github.com/sys-bio/Libstructural)

## Background

One of the most fundamental processes in living organisms is the chemical reaction where molecules combine, decompose, change configuration or exchange molecular subunits. All living systems contain large numbers of reactions forming complex reaction networks. Such networks will obey mass-conservation resulting in properties of the network that are independent of the underlying reaction kinetics. In this paper, we describe a new portable software library that provides many facilities for analyzing the topological properties of reaction networks as a result of mass-conservation.

Reactions are often depicted with reactants appearing on the left of an equation and the products on the right. Both sides are separated by an arrow indicating the positive direction of the transformation. For example, the equation:

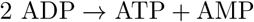

means that two molecules of ADP are transformed into one molecule of ATP and one molecule of AMP with the positive rate from left to right. The stoichiometry describes the molar amounts of reactants and products in a chemical reaction. Given a hypothetical reaction such as:

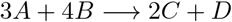

with reactants *A* and *B* and products *C* and *D*, the stoichiometry is indicated by the number of participating reactant and product molecules. Thus, the stoichiometry for *A* is three, for *B*, four, for *C*, two and for *D*, one. The number of participating molecules is sometimes referred to as the stoichiometric amounts [17].

The formal definition of the stoichiometric coefficient for a given species *X*, is the difference in stoichiometric amounts between the corresponding product and reactants. For example, given the reaction *A* → *B*, the stoichiometric coefficient for *A* is −1 because the stoichiometric amount of *A* on the product size is 0 and on the reactant side is +1, the difference is, therefore, 0 − 1 = −1. For the reaction 2*A* → *A* + *B*, the stoichiometric coefficient for *A* is −1 because 1 − 2 = −1. In the majority of cases where a given species only occurs on one side of a reaction, the stoichiometric coefficients for the reactants are the negative of the stoichiometric amounts and for the products, the positive of the stoichiometric amounts. For a reaction such as 3*A* → 2*B* + *C*, the stoichiometric coefficients are −3, +2 and +1 respectively.

When describing multiple reactions in a network, it is convenient to represent the stoichiometric coefficients in a compact form called the **stoichiometry matrix**, **N** [15]. This matrix is a m row by n column matrix where m is the number of species and n the number of reactions. The columns of the stoichiometry matrix correspond to the distinct chemical reactions in the network, the rows to the molecular species, one row per species. The intersection of a row and column in the matrix indicates the stoichiometric coefficient. For example, consider the branched pathway with two species and five reactions shown in Figure 1. The five reactions are labeled from *v*_1_ to *v*_5_ and the species labeled *S*_1_ and *S*_2_. With two species and five reactions, the stoichiometric matrix will have two rows and five columns and is given by:

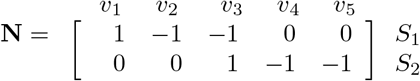

**Figure 1:**
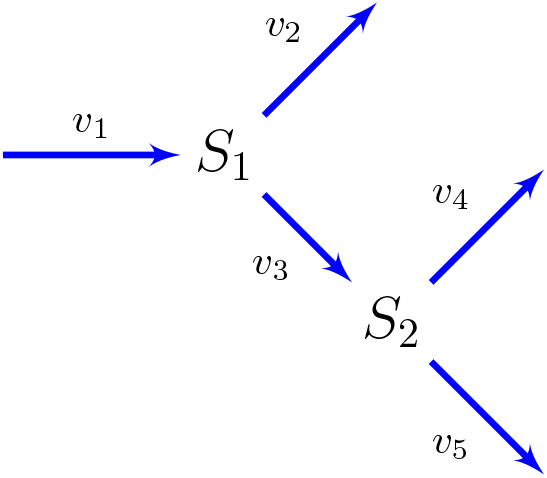
Multi-Branched Pathway with Corresponding Stoichiometry Matrix.

The stoichiometry matrix represents the connectivity of the network and contains important information on the network’s structural characteristics. These characteristics fall into two groups, relationships among the species as indicated by dependencies in the rows of the stoichiometry matrix and relationships among the reaction rates due to dependencies among the columns [21]. In this paper, we will describe a software library called `libStructural` that provides a wide variety of functions to analyze both row and column dependences in a stoichiometry matrix.

### Moiety Conservation Laws

One of the characteristics of biological network models is the conservation of certain molecular subgroups, termed moieties [16]. A typical example of a conserved group in a model is the conservation of the adenine nucleotide moiety, i.e. the total amount of ATP, ADP, and AMP is constant during the evolution of the system. It should be noted that moiety conservation is a characteristic of the model rather than the actual system itself. In practice, there will always be external inputs and outputs that render the conservation cycles open. However, in model construction it is often assumed that some processes, such as net synthesis and degradation of adenine nucleotides are slow compared to the interconversion rates, therefore over the time scale of the model study, conservation is a reasonable property to invoke.

Determining the conservation laws is important for several reasons. One practical advantage is that the system equations in the form of ordinary or stochastic differential equations can be reduced in size thus making numerical analysis more efficient. This fact is exploited in many modern modeling platforms including but not limited to SBW [20], Copasi [7], PySCeS [13], VCell [11], JWS Online [14], libRoadRunner [26], and the SB Toolbox [23]. In addition to reducing the size of the model, reduction of the number of differential equations means that the model’s Jacobian matrix is non-singular [21], an important requirement for a number of numerical methods including steady-state analysis and bifurcation analysis. The conservation laws are also important for theoretical reasons because a non-singular Jacobian is required for metabolic control analysis, stability analysis and frequency analysis [9]. Finally, conservation laws have very practical implications for perturbation studies and targeted gene knockouts [4]. In such circumstances, the conservation laws provide hard limits to how species levels can change, at least over the time scale of the conservation law. The conservation relationships can also lead to so-called implicit regulatory effects in a network as exemplified by the work of Markevich [12].

The time evolution of a biochemical network can be described by the following relation [15, 3]:

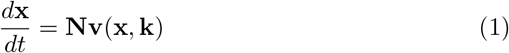

where **x** = [*x*_1_, *x*_2_,…, *x_m_*]*^T^* is the vector of time-dependent species concentrations, **N** is the stoichiometric matrix relating the species to the reactions they participate in, and **v** = [*v*_1_(**x**, **k**), *v*_2_(**x**, **k**),…, *v_n_*(**x**, **k**)]*^T^* is the vector of rates of the reactions that comprise the network, where **k** is the vector of the kinetic parameters associated with each rate law. The superscript *T* indicates the transpose and is simply a convenience for writing the vectors in row form. Dependencies between the rows of the stoichiometry matrix indicate constraints among the species of the network. Mathematically, this means that the rank, mo, of the **N** matrix is less than the number of rows in the matrix. The *m*_0_ rows correspond to independent species in the network and *m* − *m*_0_ dependent species. Let us denote these as **x_I_** and **x_D_**. The time evolution of the biochemical network described in equation (1) can then be expressed in terms of dependent and independent species as [6]:

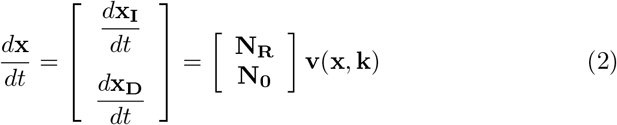

Strictly speaking **x** should be written as a function of time as in **x**(*t*), but we have dropped this notation to reduce clutter. The rows of matrix **N** have been rearranged in equation (2) such that the independent rows form a *m*_0_ × *n* matrix **N_R_** that is of full rank, and a (*m* − *m*_0_) × *n* matrix **N_0_** that comprises the dependent rows of the matrix. Equivalently, the dependent rows can be constructed as a linear combination of the independent rows. Hence a relation linking **N_R_** and **N_0_** can be written as **N_0_** = **L_0_N_R_**, where **L_0_** is a (*m* − *m*_0_) × *m*_0_ matrix. Therefore, the dynamics of the full system described by equation (2) can be rewritten as:

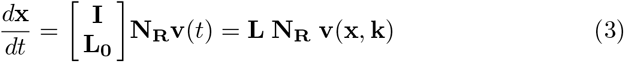

where the *m* × *m*_0_ matrix **L** in equation (3) is called the Link matrix [15]. The dynamics of the full system described by equation (1) can be partitioned into two components, one describing the dynamics of the independent species, and another corresponding to the dependent species. These can be written separately as:

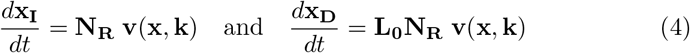

Further simplification of equation (4) yields the relation between the time evolution of the dependent species and independent species as:

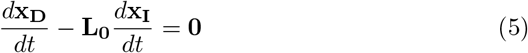

It follows that upon integrating equation (5) that **x_D_** and **x_I_** are related by a constant vector **T** = [*T*_1_, *T*_2_,…, *T*_*m−m*_0__]*^T^* such that:

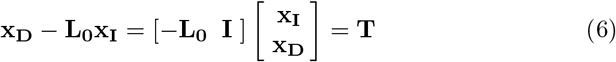

This shows that the dynamics of the dependent species can be computed directly from the independent species assuming **L**_0_ and **T** are given. In software, it is common to solve the differential equation for the independent species and to compute the dependent species using equation (6). If we replace [−**L_0_ I**] with **Γ** in equation (6), we can rewrite (6) as **Γx** = **T**. The (*m* – *m*_0_) × *n* matrix **Γ** is called the conservation matrix, because it relates the species vector **x** to the vector of conserved totals **T**. Each row represents a molecular subgroup that is conserved during the evolution of the network [16]. The values of the vector **T** can in practice be obtained by substituting the initial conditions of the species into the relation:

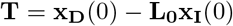

where **x_D_**(0) and **x_I_**(0) indicate the levels of **x_D_** and **x_I_** at time zero. It is important to appreciate that the conservation laws apply whether the system is at the steady-state or not. Of interest is that the relation **N_0_** = **L_0_N_R_** can be rewritten in the form:

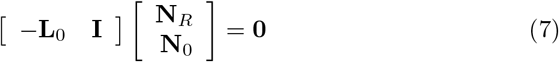

Substituting [−**L_0_ I**] with **Γ** and taking the transpose on both sides yields:

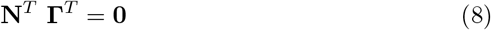

In order words, the conservation matrix can be obtained from the null space of the transpose of **N** (also called the left null space). This result provides a very quick and simple way to compute the conservation matrix in software packages such as Matlab, SciLab, or Mathematica where computing the null space of a matrix is a built-in function.

The `libStructural` library supports the computation of **L**, **L**_0_ and **Γ** matrices. In addition, the library will reorder the stoichiometry matrix rows including row labels as appropriate.

### Steady-State Flux Constraints

Whereas the rows of the stoichiometry matrix indicate dependencies among the species, the columns of the stoichiometry matrix indicate dependencies among the reaction rates. At the steady-state when all rates of change are zero, the system equation (1) reduces to:

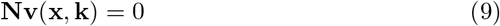

Consider a simple branched pathway with one reaction having a rate of *J*_1_ and producing a species *S*_1_, and two *S*_1_ consumption reactions with rates *J*_2_ and *J*_3_. At the steady-state it must be true that *J*_1_ = *J*_2_ + *J*_3_. Hence if we know two of the rates the third is automatically determined. In general, for an arbitrary reaction network, it is possible to divide the network rates into a set if independent and dependent sets of rates when the system is at the steady-state.

If the reaction rates are separated into a independent (**J***_I_*) and dependent (**J***_D_*) set of rates, the steady-state equation can be reexpressed as:

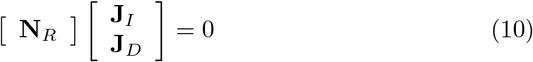

We have switched from using the symbol *v* to represent rates to use *J* instead which represents the steady-state fluxes. This helps to emphasize that the flux dependencies only apply a steady-state. **N***_R_* is the row reduced stoichiometry matrix which can, in turn, be partitioned into a set of dependent and independent columns, **N***_DC_* and **N***_IC_*.

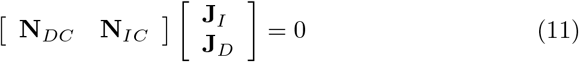

where **N***_DC_* represents the set of linearly dependent columns and **N***_IC_* the set of linearly independent columns. Multiplying out this equation gives **N***_DC_* **J***_I_* + **N***_IC_* **J***_D_* = 0. This equation can be rearranged and both sides multiplied by the inverse of **N***_IC_* to obtain:

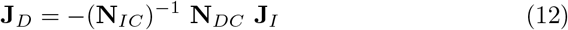

The term −(**N**_*IC*_)^−1^ **N***_DC_* can be replaced by **K**_0_ so that **J***_D_* = **K**_0_ **J***_I_*. The inverse of **N***_IC_* is guaranteed to exist because the matrix is square and all rows and columns are guaranteed by construction to be linearly independent. The equation, **K**_0_ = −(**N***_IC_*)^−1^ **N***_DC_* can be rearranged into the following form:

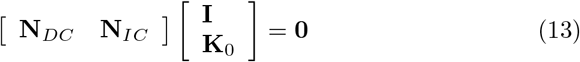

or more simply:

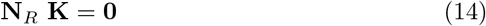

This shows that the **K** matrix is related to the null space (sometimes called the right null space) of the reordered stoichiometry matrix. The **K** matrix is important in a number of applications including relating measurable to non-measurable reaction rates, and in some algorithms as a starting point for elementary mode determination [29].

The `libStructural` library supports the computation of **K**, **N***_DC_*, and **N***_IC_* matrices. In addition, the library will reorder the stoichiometry matrix columns including column labels as appropriate. Figure 2 summarizes the partitioning of the matrix into the various partitions.

**Figure 2:**
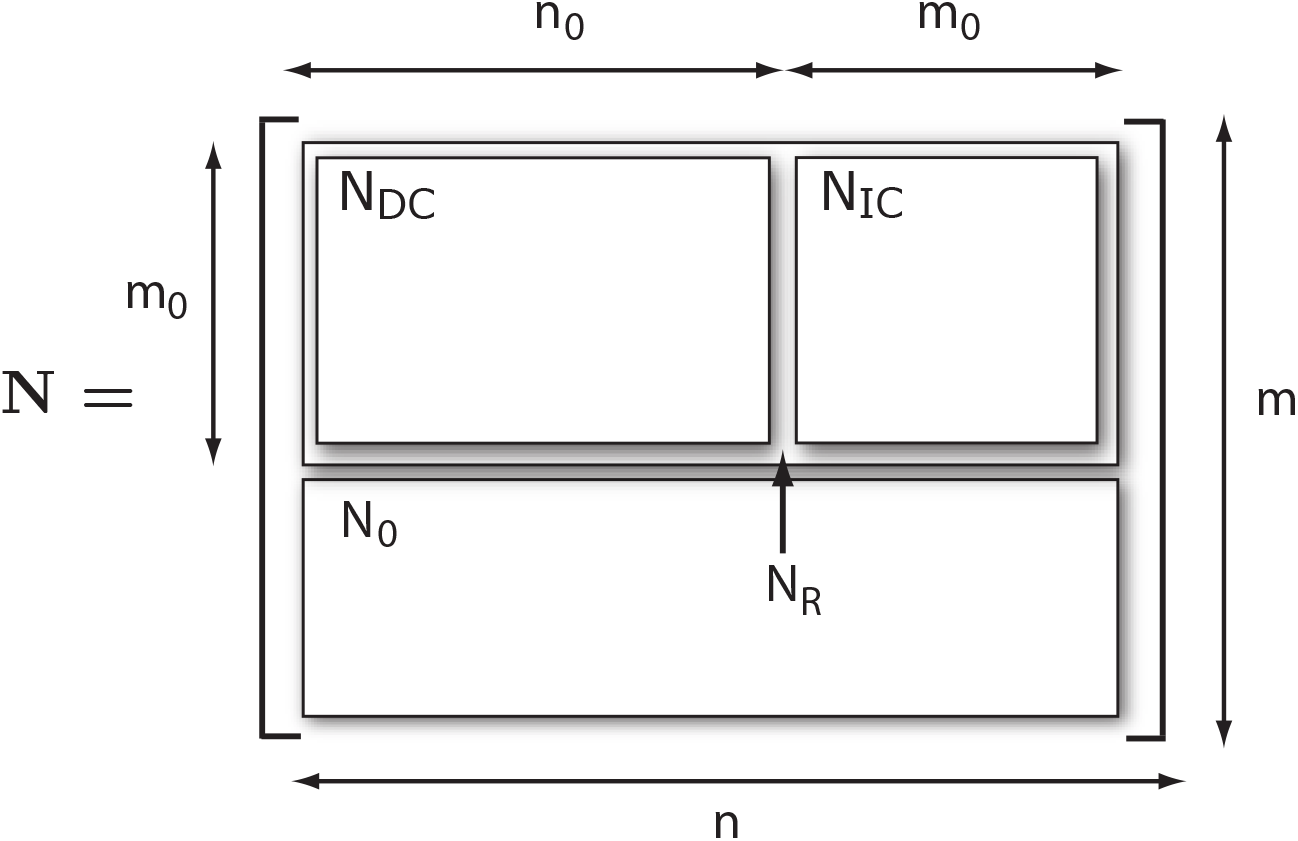
Partitioned Reordered Stoichiometry Matrix: *n* = number of reactions; *m* = number of species; **N***_DC_* = partition of linearly dependent columns; **N***_IC_* = partition of linearly independent columns; **N***_R_* = reduced stoichiometry matrix; **N**_0_ partition of linearly dependent rows.

### Elementary Modes

Elementary modes [30] are the simplest pathways within a metabolic network that can sustain a steady-state and at the same time are thermodynamically feasible. Depending on the size of the metabolic network, the number of elementary modes can range from a single mode to billions of modes. The full set of elementary modes represents the complete metabolic potential of a given metabolic network and as a result is of interest to the metabolic analysis and engineering communities.

Elementary modes are solutions to the steady-state equation but with certain constraints. An elementary mode, **e***_i_*, is defined as a vector of rates, *v*_1_, *v*_2_,…, such that the three conditions shown below must be met:

1. The vector must satisfy **Ne***_i_* = 0, which is the steady-state condition.
2. *v_i_* ≥ 0 must be true for all irreversible reactions. This means that all flow patterns must use reactions that proceed in their most natural direction. This makes the pathway described by the elementary mode a thermodynamically feasible pathway.
3. The vector **e***_i_* must be elementary, that is, it should not be possible to generate **e***_i_* by combining two other vectors that satisfy the first and second requirements using the same set of enzymes that appear as non-zero entries in **e***_i_*. In other words, it should not be possible to decompose **e***_i_* into two other pathways that can themselves sustain a steady-state.

The `libStructural` library supports the computational of elementary modes via a refactored Metatool component [10].

## Results

### Software Implementation

The core library for `libStructural` was originally developed by Frank Bergmann and was used by the original C# version of RoadRunner [1]. With the development of the C/C++ version of libRoadRunner, `libStructural` was subsequently integrated into libRoadRunner [26]. In this paper, we describe the separate and reusable `libStructural` library.

The core of `libStructural` is written in ISO C/C++ in order to achieve maximum portability and interoperability. The software can be easily used on Windows, Mac OS X, and Linux operating systems. Network models can be supplied to the software in two ways, either directly as a raw stoichiometry matrix or indirectly as an SBML model file [8]. For SBML support we use the libSBML library [2]. In order to maintain information on the row and column reorderings during the calculations, all row and columns of matrices can be labeled. Row or column exchanges are reflected in changes to the row and column labels. The library relies heavily on LAPACK (http://www.netlib.org/lapack/), a standard library for linear algebra that is used to carry out householder reflections for the QR factorization [5]. The algorithms used to carry out the computations can be found in Vallabhajosyula et al [27] and are described more fully in the methods section.

The library itself is split into two parts. One part is used to wrap and expose certain LAPACK functions and to implement other commonly used linear algebra results not directly supported by LAPACK. These include methods that can compute orthonormal null space vectors or generate the reduced echelon forms using Gauss-Jordan matrix reduction. The second part implements the stoichiometric network specific methods. These include conservation analysis such as computing the link and gamma matrices (conservation matrix), returning the reordered row stoichiometry matrix and total amounts in each conservation law. In addition, the columns of the stoichiometric matrix are also analyzed to generate the independent and dependent fluxes, including the **K** matrix. Documentation is provided through readthedocs service (https://libstructural.readthedocs.io). The source code is licensed under a combination the modified BSD license and public domain (`libMetatool`).

For computing elementary modes there are a wide variety of published software tools. Rather than write our own we decided to reuse existing software. However, it was discovered that almost all of the currently available tools are not suitable for reuse. We could not use any tools that are written in Matlab because of Matlab’s proprietary nature and limited ability to host in other applications (such as Python). We could not use any tools written in Java due to the heavy runtime requirements and the difficulty in hosting Java-based software in other applications. We did find at least one software application written in C++ but it had a restrictive usage license. There were other tools that were only available on one platform and making them cross-platform would be time consuming. Finally, there are monolithic tools such as CellNetAnalyzer [28] or COPASI where the code is tightly embedded in the application and cannot be easily extracted.

Ultimately we were left with the original elementary code application, Metatool 4.3 [10]. Interestingly this is also used by PySCeS [13], presumably for similar reasons. Metatool 4.3 has a liberal open license and because it uses standard C, it is portable across different computing platforms. The one drawback is that Metatool 4.3 is a monolithic application, therefore we had to refactor the code to convert Metatool 4.3 into a reusable library we call `libMetatool`. This allows Metatool to be linked to any programming language. `libStructural` therefore includes Metatool 4.3 but as refactored code, `libMetatool`. The refactoring was done in such a way that `libMetatool` can be used independently of `libStructural`. The one major change to Metatool when refactoring was to use long integer types for all calculations. The long integer type on a 64-bit machine is 64-bits in length and has a range of − 9, 223, 372, 036, 854, 775, 808 to 9, 223, 372, 036, 854, 775, 807. This was done to improve the numerical stability of the algorithms used by Metatool when dealing with large models. Since the original Metatool source code is in the public domain, the refactored Metatool source code is similarly unrestricted.

In addition to the C/C++ library, we also created Python bindings to allow users access to the functionality of `libStructural` via Python. Python is an easy to learn interpretive programming language that is gaining widespread usage in the scientific community. We used the SWIG toolkit (http://www.swig.org/) to generate the Python bindings and have deployed the Python enabled `libStructural` via standard pip packaging. This makes it fairly trivial to install `libStructural` on user machines.

## Applications and Examples

The following examples show how `libStructural` is used from Python for the analysis of two different metabolic networks: A complex branched pathway and a network that contains two conserved moieties.

### Branched Metabolic Network

Consider a metabolic network with nine reactions and six floating species shown in Figure 3. The model was described using the antimony syntax [25] which was then exported as SBML.

**Figure 3:**
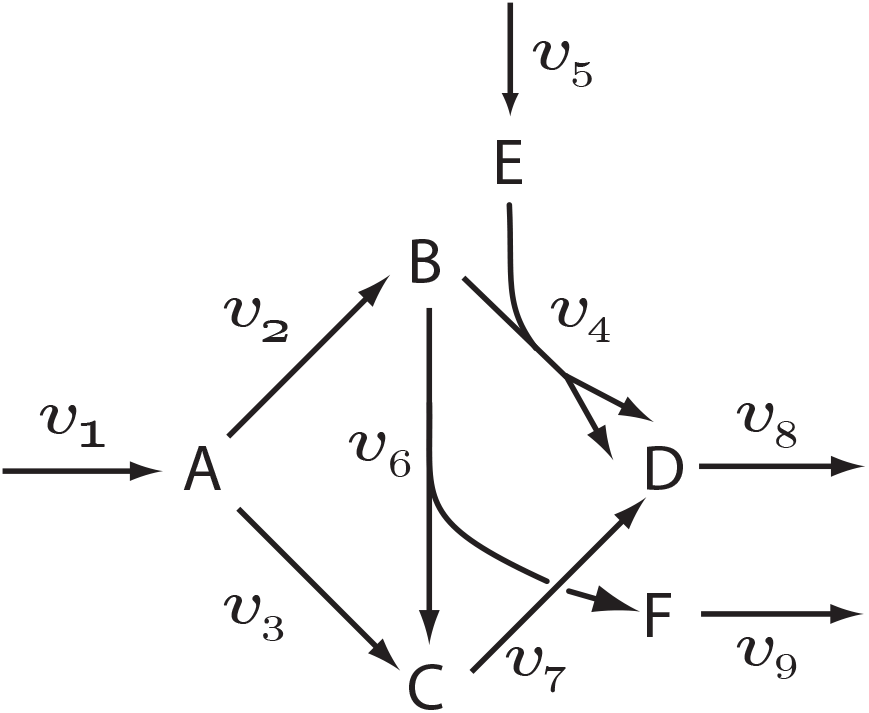
Branched network

An SBML model can be loaded by calling the `loadSBMLFromFile` function. By default, the library will apply QR factorization to the model.

~~~
>>> import structural
>>> ls = structural.Libstructural()
>>> ls.loadSBMLFromFile(“branched_network.xml”)
~~~

Models can also be conveniently loaded using the Antimony syntax [25] as shown below:

~~~
import tellurium as te
import roadrunner, structural, antimony
ant = ″″″
  *$Xo –> x; k1⋆Xo*;
  *x –> $X1; k2⋆x*;
  *Xo = 1; k1 = 0.5; k2 = 0.15*;
″″″
antimony.loadAntimonyString (ant)
ls = structural.LibStructural()
ls.loadSBMLFromString (antimony.getSBMLString(None))
print ls.getStoichiometryMatrix()
~~~

After loading a model, a summary of the analysis can be obtained using the getSummary method as shown below:

~~~
>>> ls.getSummary()
−−−−−−−−−−−−−−−−−−−−−−−−−−−−−−−−−−−−−−−−−−−−−−−−−−−−−−−−−−−−−−
STRUCTURAL ANALYSIS MODULE : Results
−−−−−−−−−−−−−−−−−−−−−−−−−−−−−−−−−−−−−−−−−−−−−−−−−−−−−−−−−−−−−−
Size of Stochiometric Matrix: 6 × 9 (Rank is 6)
Nonzero entries in Stochiometric Matrix: 16 (29.6296% full)
Independent Species (6) :
D, A, C, F, E, B
Dependent Species : NONE
L0 : There are no dependencies. L0 is an EMPTY matrix
Conserved Entities: NONE
~~~

Specific information can be obtained by calling a variety of methods. For example, the stoichiometry matrix is obtained as follows:

~~~
>>>ls.getStoichiometryMatrix()
array([[  1.,  −1., −1.,  0., 0., 0.,  0., 0., 0.],
[ 0., 1., 0., −1., 0., −1.,  0.,  0.,  0.],
[ 0., 0., 1.,  0., 0.,  1., −1.,  0.,  0.],
[ 0., 0., 0., −1., 1.,  0.,  0.,  0.,  0.],
[ 0., 0., 0.,  2., 0.,  0.,  1., −1.,  0.],
[ 0., 0., 0.,  0., 0.,  1.,  0.,  0., −1.]])
~~~

The order of species and reactions in the rows and columns can be obtained using the methods `getFloatingSpeciesIds` and `getReactionIds`. At this stage, neither the rows or columns have been reordered into dependent and independent rows and columns. A call to `getFullyReorderedStoichiometryMatrix` will return a stoichiometry matrix where both rows and columns have been reordered:

~~~
>>>ls.getFullyReorderedStoichiometryMatrix()
array([[ 1., −1., 0., 2., 0., 0.,  0., 0., 0.],
[ 0., 0., −1.,  0.,  0., −1., 0.,  0., 1.],
[−1., 0.,  0.,  0.,  1.,  1., 0.,  0., 0.],
[ 0., 0.,  0.,  0.,  1.,  0., 0., −1., 0.],
[ 0., 0.,  0., −1.,  0.,  0., 1.,  0., 0.],
[ 0., 0.,  1., −1., −1.,  0., 0.,  0., 0.]])
~~~

The ordering of the rows and columns can be obtained by calling `getFully-ReorderedStoichiometryMatrixIds`. This returns two lists, the reordered rows labels and the reordered column labels. Reordering of only the rows can be obtained by calling `getReorderedStoichiometryMatrix`.

Of particular interest are the dependent and independent reaction rates, obtained by calling `etNumDepReactions` and `getNumIndReactions`. These functions will return the number of dependent and independent reactions in the model. In the case of the example model, these methods return 3 and 6 respectively. `getDependentReactionIds` and `getIndependentReactionIds` will return the Ids of the specific reactions in each group. For example:

~~~
ls.getDependentReactionIds()
Out[24]: (‘J7’, ‘J8’, ‘ J2’)
>>>ls.getIndependentReactionIds()
Out[25]: (‘J4’, ‘J6’, ‘J3’, ‘J5’, ‘J9’, ‘J1’)
~~~

In practical terms this means that given the fluxes through (‘J4’, ‘J6’, ‘J3’, ‘J5’, ‘J9’, ‘J1’) it is possible to determine the remaining three fluxes: (‘J7’, ‘J8’, ‘J2’). A full range of methods are available to extract any of the submatrices shown in Figure 2.

### Network Containing Conserved Moieties

The model shown in Figure 4 illustrates a simple reaction network that contains two conserved moieties, namely S and E. These result in two conservation laws: *S*_1_ + *S*_2_ + *ES* and *ES* + *E*.

**Figure 4:**
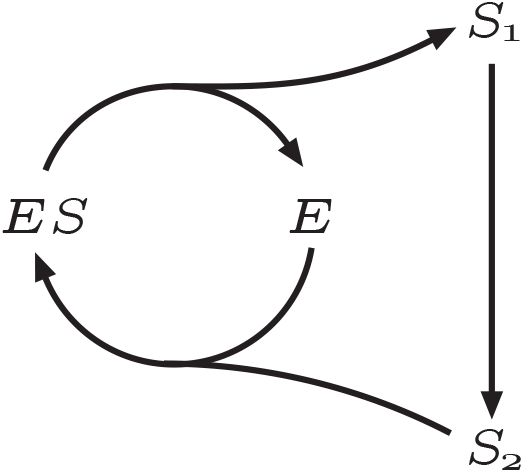
Network with Two Interlinked Conserved Moieties.

After loading the model, calling getSummary generates the following output:

~~~
>>> ls.getSummary()
−−−−−−−−−−−−−−−−−−−−−−−−−−−−−−−−−−−−−−−−−−−−−−−−−−−−−−−−−−−−−−
STRUCTURAL ANALYSIS MODULE : Results
−−−−−−−−−−−−−−−−−−−−−−−−−−−−−−−−−−−−−−−−−−−−−−−−−−−−−−−−−−−−−−
Size of Stochiometric Matrix: 4 × 3 (Rank is 2)
Nonzero entries in Stochiometric Matrix: 8 (66.6667% full)
Independent Species (2) :
ES, S1
Dependent Species (2) :
E, S2
L0 : There are 2 dependencies. L0 is a 2×2 matrix.
Conserved Entities
1: + ES + E
2: + ES + S1 + S2
~~~

From the summary, we can see that there are two dependent rows (species) and two conserved entities. Thus, the conservation matrix (gamma, **Γ**) will have two rows.

~~~
>>>ls.getStoichiometryMatrix()
array([[−1., 0., 1.],
       [ 1., −1., 0.],
       [1., 0., −1.],
       [0., 1., −1.]])
>>> ls.getGammaMatrix()
array([[ 1., 0., 1., 0.],
       [1., 1., 0., 1.]])
~~~

In addition, it is possible to obtain the conserved sums and conserved laws of the network by calling `getConservedLaws` and `getConservedSums` respectively.

~~~
>>>ls.getConservedLaws()
(‘ + ES + E’, ‘ + ES + S1 + S2’)
>>>ls.getInitialConditions()
((‘ES’, 10.0), (‘S1’, 10.0), (‘E’, 10.0), (‘S2’, 10.0))
>>> ls.getConservedSums()
(20.0, 30.0)
~~~

The function `getFullyReorderedStoichiometryMatrix` divides the **N***_R_* matrix into **N***_IC_* and **N***_DC_* matrices as shown in Fig 2.

~~~
>>> ls.getFullyReorderedStoichiometryMatrix()
array([[ 1., −1., 0.],
       [0., 1., −1.],
       [−1., 1., 0.],
       [−1., 0., 1.]])
>>>ls.getFullyReorderedNrMatrix()
array([[ 1., −1., 0.],
       [0., 1., −1.]])
>>> ls.getNICMatrix()
array([[−1., 0.],
       [ 1., −1.]])
>>> ls.getNDCMatrix()
array([[ 1.],
       [ 0.]])
~~~

### Calculating elementary modes

To illustrate the evaluation of the elementary modes, consider the previous branched model in Figure 3. The representation of this model using Antimony syntax makes it easy to specify whether a given reaction is reversible or not. Reactions that are irreversible are indicated by = while reactions that are irreversible are indicated with =>.

~~~
J1: $Xo => A; v;  J2: A => B; v;
J3: A => C; v;           J4: B + E => 2 D; v;
J5: $X1 => E; v;  J6: B => C + F; v;
J7: C => D; v;           J8: D => $X2; v;
J9: F => $X3; v;  v = 0;
~~~

If all reactions are irreversible, then the model admits only three elementary models. If all reactions are reversible then ten elementary models are possible (not shown). From the Antimony script, we can generate standard SBML which, as before, can be loaded into `libStructural`. Calling the getElementary-Modes function returns the following three elementary modes.

~~~
>>> ls.getElementaryModes()
[[ 1. 1. 0. 1. 1. 0. 0. 2. 0.]
 [ 1. 0. 1. 0. 0. 0. 1. 1. 0.]
 [ 1. 1. 0. 0. 0. 1. 1. 1. 1.]]
~~~

The elementary modes in the output are arranged in rows. The order of the columns in the output can be obtained by calling `getReactionIds` which returns the list of reaction names. To assist in visualizing the three modes, Figure 5 highlights the reactions involved in each of the modes. This shows there are three independent and thermodynamically feasible pathways in the network: 1) *v*_1_ to *v*_9_ via *v*_2_ and *v*_4_; 2) *v*_1_ to *v*_8_ via *v*_2_ and **v*_6_;* and 3) *v*_1_ to *v*_9_ via **v*_3_* and *v*_7_. All possible thermodynamically feasible flux patterns can be obtained by linear combinations of these three.

**Figure 5:**
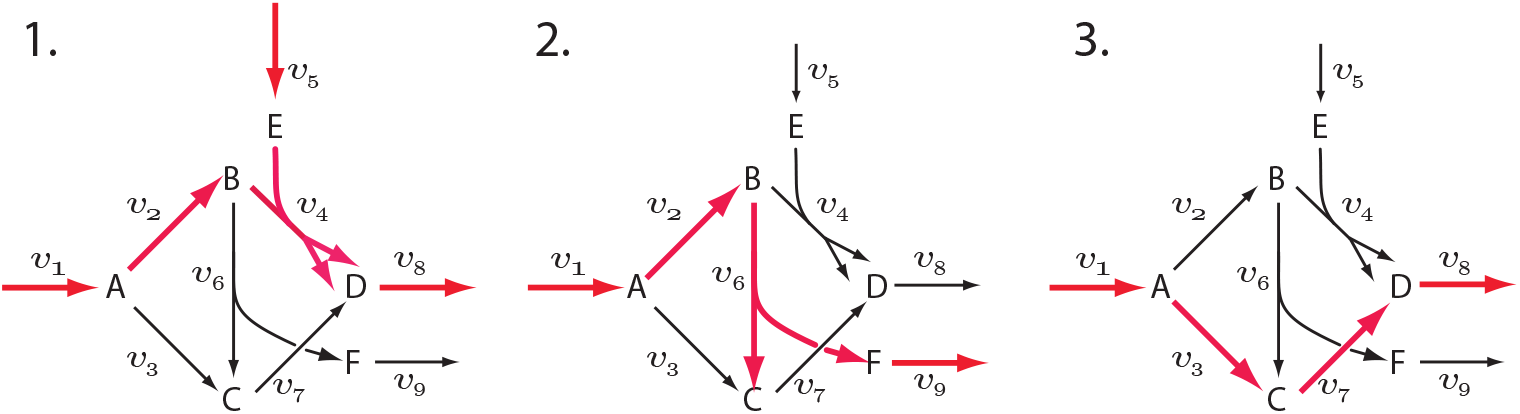
Three Elementary Modes in a Complex Pathway.

### Plugin for PathwayDesigner

PathwayDesigner is a Microsoft Windows-based reaction network editor and viewer (Available at http://pathwaydesigner.org). The application has a plugin architecture that allows new functionality to be added via external tools. PathwayDesigner exposes an application programming interface (API) to plugins that allow them to interact with the visual representation of the network. An obvious use case is to allow an external plugin to manipulate or highlight particular aspects of a network. In this case, we can compute the elementary models and display the modes on the visual representation of the network.

Two plugins were implemented for PathwayDesigner. The first was a non-visual plugin that provides a programmatic interface to the `libStructural` library. A second plugin was a visual plugin that provides a programmatic link between PathwayDesigner itself and the `libStructural` plugin. The visual plugin can load a model from PathwayDesigner and offer a number of options to the user, including obtaining the stoichiometry matrix, the conservation laws, and the elementary modes. Figure 6 shows PathwayDesigner with the plugin window visible and one of the elementary modes displayed on the PathwayDesigner canvas. Modes are highlighted using a thicker reaction edge and user-specific color (Figure 6). Users can move from one elementary mode to the next by selecting a mode from the list of modes.

**Figure 6:**
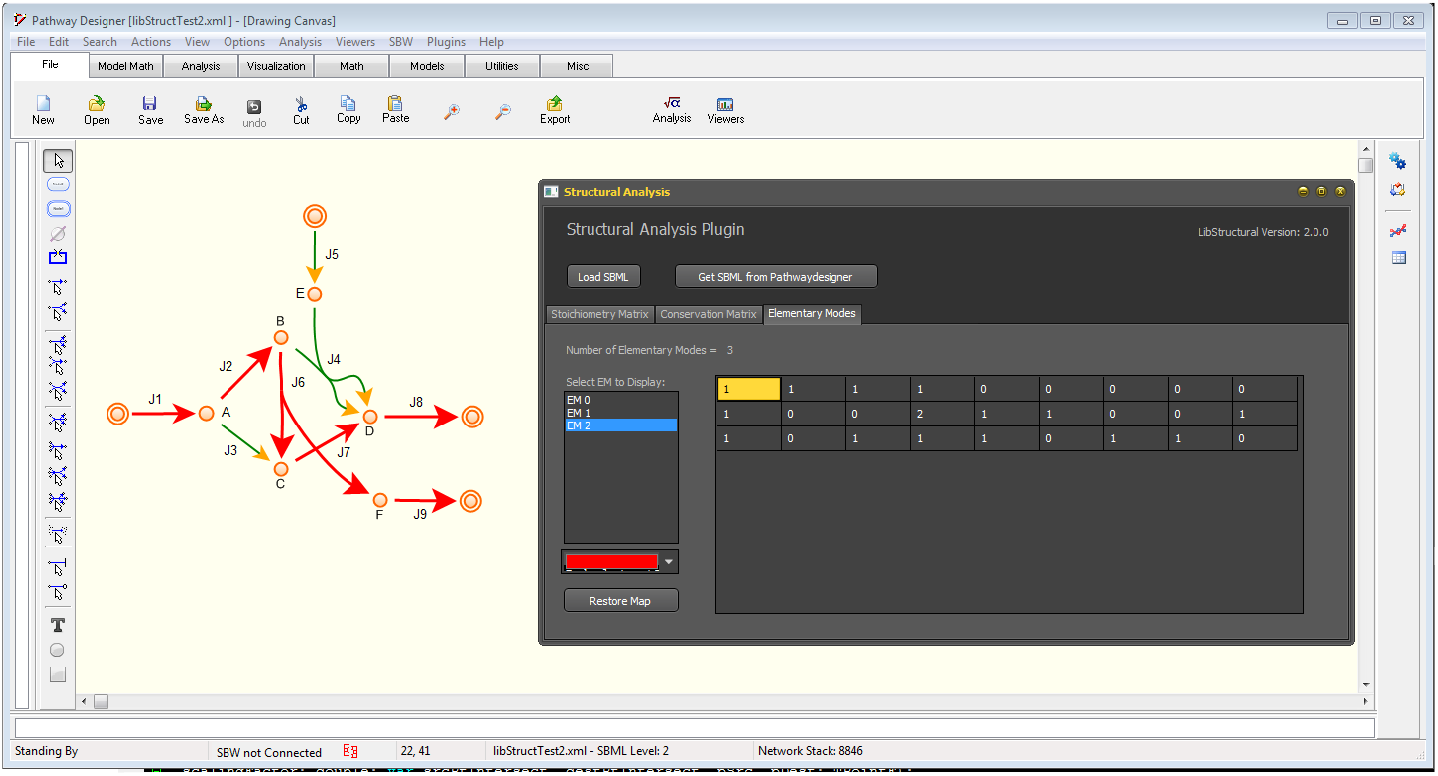
Pathwaydesigner Illustrating libStructural Plugin Displaying Elementary mode.

### Availability

`LibStructural` is available either as source code or as precompiled modules for Python. The source code is available on GitHub (https://github.com/sys-bio/Libstructural). CMake (https://cmake.org/) configurations are supported to generate build systems for different compilers and platforms. SWIG (http://www.swig.org/) is necessary for generating Python language support. Extensive instructions on how to build `libStructural` on Microsoft Windows and Apple Mac OS X are available on https://libstructural.readthedocs.io.

For ease of use, `libStructural` can be installed in Python via pip, the Python package manager. Support for Python 2.7 and 3.4+ is available and standard Python users should execute `pip install libStructural` at the command line. When using Tellurium [19], `libStructural` can be installed directly from the python console via the installPackage function.

## Methods

Details of the algorithms can be found in [24, 27]. Here we will describe the selftest methods that have been included in `libStructural`. Much of the analysis done in `libStructural` is based Householder reflections [18] to compute the QR factorization of the stoichiometry matrix. The QR decomposition results in a very robust numerical scheme to extract the species and flux dependencies. We illustrate the approach using a number of large models obtained from the BiGG repository [22] at the University of San Diego, California (see http://bigg.ucsd.edu//). The results of these analyzes are shown in Table 1.

**Table 1:** Results of Analyzing a Selection of Large Models from http://bigg.ucsd.edu/. Dep Rxns indicates the number of dependent reactions and Ind Rxns the number of independent reactions. sp is short-hand for species.

| Model | # Conserved Moieties | # Dep Rxns | # Ind Rxns |
| --- | --- | --- | --- |
| RECON1<br>(3741 rx, 2766 sp.) | 92 | 1067 | 2674 |
| IJR904<br>(1975 rx, 761 sp) | 18 | 332 | 743 |
| IND750<br>(1266 rxn, 1059 sp) | 77 | 284 | 982 |
| iJO1366<br>(2583 rxn, 1805 sp) | 39 | 817 | 1766 |
| iNRG857_1313<br>(2735 rxn, 1311 sp) | 60 | 852 | 1883 |
| iYO844<br>(1250 rxn, 990 sp) | 31 | 291 | 959 |
| iYL1228<br>(2262 rxn, 1658 sp) | 45 | 649 | 1613 |
| STM_v1_0<br>(2545 rxn, 1802 sp) | 39 | 782 | 1763 |

### Self-Test Routines

The conservation laws generated by the algorithm must be tested to ensure that they are valid. Therefore, the library includes five tests to validate the conservation laws. These tests need only be carried out for very large models. The self tests can be activate by calling validateStructuralMatrices() which returns a list of a true/false values or getTestSummary() which returns a string giving a more detailed account, for example, getTestSummary() will yield:

~~~
Passed Test 1 : Gamma⋆N = 0 (Zero matrix)
Passed Test 2 : Rank(N) using SVD (1) is same as m0 (1)
Passed Test 3 : Rank(NR) using SVD (1) is same as m0 (1)
Passed Test 4 : Rank(NR) using QR (1) is same as m0 (1)
Passed Test 5 : L0 obtained with QR matches Q21⋆inv(Q11)
Passed Test 6 : N⋆K = 0 (Zero matrix)
~~~

The first test checks if the conserved sums are indeed constant values. This is done by observing that the conserved sums are given by **T** = **ΓS**. Their rate of change, *d***T**/*dt* must be zero if **T** is a constant vector. This is equivalent to

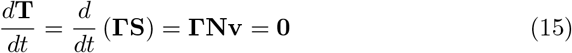

The second test checks if the rank obtained by QR factorization of the stoichiometry matrix **N** is the same as that obtained by counting the non-zero singular values obtained from an SVD of **N**. If the rank computed by the two methods is different, it is evident that the number of conserved cycles has been miscalculated.

The third test checks the rank of **N_R_** matrix. Since the **N_R_** matrix corresponds to the contribution of all the independent species in the network, it must have a full rank. Therefore, one can compare the rank of **N_R_** using QR factorization to that obtained by SVD. If these ranks are full and match, then the conservation laws are correct.

The fourth test involves the computation of the eigenvalues of a sub-matrix obtained from the QR factorization of **N**. We note that the orthogonal matrix **Q** obtained after factorizing the reordered stoichiometric matrix can be partitioned into the following sub-matrices as described by:

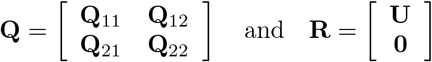

If the stoichiometric matrix **N** has been reordered such that the first *m*_0_ rows correspond to independent species and the remaining (*m* − *m*_0_) rows correspond to dependent species, it can be shown that **Q**_11_ is a non-singular and invertible matrix. Therefore, we can compute its rank by counting the number of nonzero eigenvalues of **Q**_11_. If **Q**_11_ is of full rank, that is if *m*_0_ = *rank*(**Q**_11_), the conservation laws must be correct.

The fifth test evaluates the **L**_0_ matrix as the product of the sub-matrices of **Q**. As shown in [27], 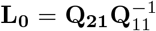. The matrix obtained by multiplying **Q**_21_ and the inverse of **Q**_11_ is compared with the **L**_0_ matrix obtained directly by the Householder method. The test can then be considered successful if both matrices match. Similar tests can be done when computing the **K** matrix. However, passing the conservation and rank tests is sufficiently stringent to also validate simultaneously the flux dependencies and the **K** matrix.

The results of these validity tests depend on the size of the model and the method employed to generate the conservation laws. The numerical errors inherent in the algorithm are highest in the case of LU decomposition with partial pivoting, leading to the generation of conservation laws that do not pass the validity tests. On the other hand, the Householder QR method used in `libStructural` and LU decomposition with full pivoting (used by PySCeS) generate the same number of conservation laws, both of which pass the validity tests.

There is one caveat to consider. Due to rounding errors during the computation, it is important to be able to distinguish between zero and small numbers. This is accomplished by setting a tolerance in the library (`setTolerance`). By default the tolerance is set to 10^−9^, that is any number less than 10^−9^ is considered to be zero. For large models, it may be necessary to reduce the tolerance to 10^−6^ in order to pass the tests. For extremely large models of the order of 100,000 nodes, new approaches may be necessary. However, the built-in tests can be used to determine whether the results from large studies are correct or not.

### Tests for Elementary Modes

There are three tests that can be done to check that the elementary modes, e, enumerated by Metatool are elementary modes. Due to the current state of understanding of elementary models, it is unfortunately not possible to determine before-hand how many elementary modes to expect. Therefore it is not possible to know with certainty whether all elementary modes have been identified. This is particularly true for larger networks where the number of elementary modes can be extremely large. The tests we have used to test Metatool include the following:

1. All elementary modes must satisfy **Ne***_i_* = 0.
2. For a given elementary mode, **e***_i_*, it must be true that:

a. The null space of the submatrix of **N** that only involves the reactions of **e***_i_* is of dimension one and has no zero entries (elementarity).
b. The elementary mode must be consistent with respect to the sign of the coefficient for irreversible reactions (thermodynamic correctness). That is, if a reaction is irreversible, its corresponding sign in **e***_i_* must be positive. A reversible reaction may be negative or positive.

The test file `elementaryModes.py` includes 21 models and each model is tested against the above tests.

## Authors contributions

HMS conceived the idea and FTB implemented the methods, testing, and documentation. YB build the distributions, the Python bindings, documentation, and testing. KC advised YB on builds and bindings and help create the CMake files. Everyone contributed to the manuscript.

## Acknowledgements

Much of the original work described in this paper is due to the generous support from the NIH grant GM081070 and more recently from GM123032. We would also like to thank David Fell and Mark Poolman for constructive discussions regarding the tests for elementary modes.

